# Dynamics of transposable elements in recently diverged fungal pathogens: lineage-specific transposable element content and efficiency of genome defences

**DOI:** 10.1101/2020.05.13.092635

**Authors:** Cécile Lorrain, Alice Feurtey, Mareike Möller, Janine Haueisen, Eva Stukenbrock

## Abstract

Transposable elements (TEs) impact genome plasticity, architecture and evolution in fungal plant pathogens. The wide range of TE content observed in fungal genomes reflects diverse efficacy of host-genome defence mechanisms that can counter-balance TE expansion and spread. Closely related species can harbour drastically different TE repertoires, suggesting variation in the efficacy of genome defences. The evolution of fungal effectors, which are crucial determinants of pathogenicity, has been linked to the activity of TEs in pathogen genomes. Here we describe how TEs have shaped genome evolution of the fungal wheat pathogen *Zymoseptoria tritici* and four closely related species. We compared *de novo* TE annotations and Repeat-Induced Point mutation signatures in thirteen genomes from the *Zymoseptoria* species-complex. Then, we assessed the relative insertion ages of TEs using a comparative genomics approach. Finally, we explored the impact of TE insertions on genome architecture and plasticity. The thirteen genomes of *Zymoseptoria* species reflect different TE dynamics with a majority of recent insertions. TEs associate with distinct genome compartments in all *Zymoseptoria* species, including chromosomal rearrangements, genes showing presence/absence variation and effectors. European *Z. tritici* isolates have reduced signatures of Repeat-Induced Point mutations compared to Iranian isolates and closely related species. Our study supports the hypothesis that ongoing but moderate TE mobility in *Zymoseptoria* species shapes pathogen genome evolution.

## Introduction

Transposable elements (TEs), DNA elements that can replicate through transposition (i.e. independently of the host genome replication machinery) are ubiquitous in Eukaryotic genomes. The TE content in fungal plant-pathogen genomes covers a wide range: from less than 1% to more than 90% of the genome in *Fusarium graminearum* and *Blumeria graminis*, respectively (Cuomo *et al*. 2007; Frantzeskakis *et al*. 2018). TEs are categorized into two classes, retrotransposons (class I) and DNA transposons (class II), based on their mechanism of transposition (Wicker *et al*. 2007). Retrotransposons replicate using an RNA intermediate to insert at a new position and DNA transposons replicate either by a mechanism of direct excision from double-stranded DNA (subclass I) or using single-strand excision followed by a rolling-circle mechanism (subclass II) (Wicker *et al*. 2007). TE classes are divided into orders that contain various numbers of superfamilies and families, which are categorized by coding sequence structure (Wicker *et al*. 2007). TEs can be autonomous (e.g. LTRs and TIRs) or non-autonomous (e.g. SINEs and MITEs). The latter relies on the replication machinery of autonomous TEs to transpose (Wicker *et al*. 2007). TE activity (i.e. transposition) is known to have an overall negative impact on host fitness (Horváth *et al*. 2017). As a result, TEs engage in a co-evolutionary arms race dynamic with the host genome (Biémont 2010; Castanera *et al*. 2016).

Fungal genomes have evolved diverse genome defence mechanisms to regulate TE expansions. In addition to histone modifications and DNA methylation (Deniz *et al*. 2019), a fungal-specific mechanism called Repeat-Induced Point mutation (RIP) specifically mutates duplicated sequences such as TEs (Gladyshev 2017). RIP induces a dinucleotide bias in duplicated sequences by mutating G:C into A:T; this bias can be measured in fungal genomes, and quantified as a RIP signature (Gladyshev and Kleckner 2014, 2017). In “RIPed” genomes, RIP-induced mutation can result in Large RIP Affected Regions (LRARs) that are large genomic regions consecutively affected by RIP (van Wyk *et al*. 2019). Genomes of several fungal pathogens have relatively high TE contents, while simultaneously exhibiting signatures of RIP (Gao *et al*. 2011; Gioti *et al*. 2013; Grandaubert *et al*. 2014; Dhillon *et al*. 2014; Fokkens *et al*. 2018). It remains unclear how TEs can maintain stable proportions in their host genomes with defence mechanisms such as RIP. It is worth to note that the maintenance of RIP mechanisms is costly for the host and RIP can be lost (Galagan and Selker 2004). In the genus *Neurospora*, closely related species have variation in RIP signatures, and *Neurospora* species with reduced RIP signatures exhibit TE expansions (Gioti *et al*. 2013). This dynamic between TE expansions and host defense mechanisms is summarized as the burst and decay model of TE evolution (Arkhipova 2018). This model assumes that TEs are active (i.e., they burst in transposition or expansion) until they are inactivated by host defence mechanisms such as RIP (i.e., they decay). As a consequence of the RIP-induced overaccumulation of mutations in TEs, dating specific TE families’ invasions is challenging (Grandaubert *et al*. 2014).

TE insertions and how the host respond to them, shape genome architecture. TEs can increase genomic plasticity by promoting chromosomal rearrangements and compartmentalization, duplicating or deleting genes, and altering gene expression. Often, the TE content is linked to compartmentalization of the genome with TE-enriched genomic compartments or regions associating with specific epigenetic signatures and changes in CG content (Duplessis *et al*. 2011; Bertazzoni *et al*. 2018; Frantzeskakis *et al*. 2018; Chen *et al*. 2018; Stam *et al*. 2018). Genome compartmentalization into TE-rich and TE-poor regions can sometimes be observed at the chromosomal level. Some fungal species harbor dispensable or accessory chromosomes, which contain a higher proportion of TEs than the core chromosomes as observed in the wheat pathogen *Zymoseptoria tritici* (Croll and McDonald 2012). Despite being considered as “junk DNA” until recently, TEs have been shown in recent years to participate in adaptation to environmental changes of their hosts. For instance, in plant-associated fungi, TEs are de-repressed during stressful conditions such as the early stage of plant infection (Fouché *et al*. 2020). TEs are also physically associated with pathogenicity-related genes (i.e., effectors) of various fungal pathogen species, suggesting a role in effector gene diversification (Rouxel *et al*. 2011; Grandaubert *et al*. 2014; Soyer *et al*. 2014; Dong *et al*. 2015; Fokkens *et al*. 2018; Fouché *et al*. 2018). In this way TEs can have mediated mutational changes with benefits to the host organism. TEs can even be co-opted, or domesticated by the host, and evolved to have a new function in the host. For instance, the effector AvrK1 in *Blumeria graminis* f.sp. *hordei* is directly derived from a LINE retrotransposon (Amselem *et al*. 2015b). In *Z. tritici*, LTR retrotransposon insertion upstream of a multidrug efflux transporter has conferred fungicide resistance (Omrane *et al*. 2017b). Multiple studies demonstrate the importance of TEs for fungal pathogenicity, yet little is known about the extent to which TEs represent key players in the global evolution of the genomes of fungal pathogens.

The wheat pathogen *Z. tritici* has emerged as a model to study fungal genome and TE evolution. Closely related species of *Z. tritici* have been isolated from wild grasses and barley, and include *Zymoseptoria passerinii*, *Zymoseptoria ardabiliae*, *Zymoseptoria brevis* and *Zymoseptoria pseudotritici* (Stukenbrock *et al*. 2007, 2012). The genomes of the five species comprise conserved core chromosomes and variable accessory chromosomes which show low gene density, low transcriptional activity, and enrichment of the heterochromatin-associated histone mark H3K27me3 (Goodwin *et al*. 2011; Kellner *et al*. 2014; Schotanus *et al*. 2015; Feurtey *et al*. 2019). TEs have been investigated in the economically important wheat pathogen *Z. tritici*, describing for example the occurrence of RIP in this species through the quantification of C to T transition mutations for the reference isolate IPO323 (Dhillon *et al*. 2014). Several associations have been made between TEs and *Z. tritici* virulence and fungicide resistance. *Z. tritici* genes encoding effectors and transporters involved in fungicide resistance physically associate with TEs (Omrane *et al*. 2017b; Hartmann *et al*. 2018; Oggenfuss *et al*. 2020). Recent studies demonstrated that TEs in *Z. tritici* are de-repressed during early stages of wheat infection (Fouché *et al*. 2020). Interestingly, the TE content varies between closely related *Zymoseptoria* species(Grandaubert *et al*. 2015). TEs in *Z. tritici* accumulate in recently founded populations outside of the center of origin due to relaxed purifying selection (Fouché *et al*. 2020; Oggenfuss *et al*. 2020; Badet *et al*. 2020). Despite these recent studies, little is yet known about how TEs and genome defence mechanisms have co-evolved and shaped host genomes over larger time scales among the *Zymoseptoria* genus.

In this study, we explore TE dynamics in thirteen *Zymoseptoria* genomes, including nine *Z. tritici* genomes and one genome from each of the four closely related species, *Z. passerinii, Z. ardabiliae, Z. brevis* and *Z. pseudotritici*. We specifically addressed the following questions: i) How do TE distributions and insertion ages impact genome architecture and plasticity? ii) Does TE content correlate with gene presence/absence variation among genomes? iii) What is the extent of variation in RIP among genomes? To answer these questions, we annotated the TE content of each of the thirteen genomes *de novo* and analysed TE landscapes within and among genomes. We found evidence for variation in the efficiency of RIP among *Z. tritici* genomes and among the different species genomes.

## Material and Methods

### Genomic and gene expression data

All *Zymoseptoria* spp. isolates used in this study come from publicly available genome assemblies. We used recently assembled genomes of nine *Z. tritici* isolates sampled from Europe and Iran. Zt05 was isolated from wheat in Denmark, Zt10 and Zt289 from wheat in Iran and Zt469 from *Aegilops* sp. in Iran (Grandaubert *et al*. 2015; Feurtey *et al*. 2019; Möller *et al*. 2020). We also included the genomes of five *Z. tritici* isolates for which recent Pacbio assemblies were published (four from Switzerland Zt1A5; Zt1E4; Zt3D1; Zt3D7 (Plissonneau *et al*. 2018). Finally, we used one genome from each of four closely related species *Z. ardabiliae* Za17, *Z. brevis* Zb87; *Z. passerinii* Zpa63 and *Z. pseudotritici* Zp13 (Feurtey *et al*. 2019). Gene expression during wheat infection from (Haueisen *et al*. 2019), updated expression profiles on the last versions of genome assemblies and new gene predictions of the three *Z. tritici* isolates IPO323, Zt05 and Zt10, were used as described in (Feurtey *et al*. 2019). In summary, RNAseq was performed during wheat infection time-course using strand-specific RNA-libraries from Illumina HISeq2500 sequencing, with 100pb single-end reads. A total of 89.5 to 147.5 million reads per sample was obtained (Haueisen *et al*. 2019). To simplify the expression data, we combined all time-points from wheat infection and calculated expression levels in Transcripts Per Million (TPM) as described in (Feurtey *et al*. 2019).

### Annotation of repeated elements and relative age of insertion

We used the REPET pipeline (https://urgi.versailles.inra.fr/Tools/REPET; (Quesneville *et al*. 2005; Flutre *et al*. 2011) to annotate the repeat regions of *Z. ardabiliae* Za17, *Z. brevis* Zb87, *Z. pseudotritici* Zp13, *Z. passerinii* Zpa63 and the nine *Z. tritici* isolates as described in (Feurtey *et al*. 2019). Briefly, we identified repeats in each genome by building TE consensus sequences, as a proxy for TE ancestral sequence. Each consensus is derived from a multiple alignment of TEs in clusters. We then mined each *Zymoseptoria* spp. genome using the constructed TE consensus library to recover TE copies belonging the same consensus. TE annotation metrics are summarized in Table S1.

We assessed relative ages of TE insertions in *Zymoseptoria* spp. using the REPET similarity-based approach to measure the distributions of TE clusters’ sequence identities. Based on the burst and decay evolution of TEs (Roessler *et al*. 2018), we analysed sequence divergence of individual element clusters to assess the relative age of TE insertions in each genome (i.e. TE spread in host genome). For this, we need to assume that RIP-induced mutations occurrence is constant over time for each genome. Based on this assumption, the extent of sequence similarity is proportional to the divergence time of copies. It is thereby possible to compare relative insertion ages of TE insertions within genomes (Figure S1). We assessed sequence similarity within each TE cluster by comparing each TE copy to the consensus sequence.

### Analysis of transposon genomic environments

To further investigate intra-specific variation in TE content, we conducted a detailed comparison of the core chromosomes of the nine *Z. tritici* isolates. By focusing only on the core chromosomes, we avoid an overestimation of TE insertions variation due to presence/absence polymorphisms of the accessory chromosomes. Transposon densities along *Z. tritici* isolates’ chromosomes were measured using 100 kbp windows using bedtools “makewindows”. We fixed window coordinates based on the reference genome of IPO323 using orthologous genes. The closest genes neighbouring each window’s borders were extracted from the reference IPO323 genome using the bedtools “closest” function (Quinlan and Hall 2010). Orthologs of these neighbouring genes were then extracted after identification using the program Proteinortho (Lechner *et al*. 2011). The number of transposons per window was calculated using bedtools “intersect” function (Quinlan and Hall 2010). The results were visualized using the software Circos (Krzywinski *et al*. 2009).

To assess associations between the vicinity of TEs and genes potentially involved in plant infection, we used the previous functional annotations of predicted effectors and orthologous genes of Za17, Zpa63, Zb87 and Zp13, and the three *Z. tritici* isolates IPO323, Zt05 and Zt10 (Feurtey *et al*. 2019). We annotated the genes with presence/absence variation (PAV) among the thirteen *Zymospetoria* species-complex genomes. For this we used PoFF, an extension of the software Proteinortho which integrates data on conserved synteny to detect orthologous relationships (Lechner *et al*. 2011). We differentiated genes as follows: 1) showing PAV among all thirteen genomes as *PAV genes*, 2) genes present in all thirteen *Zymoseptoria* genomes as *Core genes* and 3) genes present on all nine *Z. tritici* isolates as *Core Z. tritici genes*. We also included predicted genes with TE-like domains (e.g. transposase) in the TE repertoires to avoid considering TEs as PAV genes. To statistically assess associations between the vicinity of transposons and gene categories, we used the R package regioneR (Gel *et al*. 2015). We used the function “meanDistance” to test whether specific gene categories are closer to transposons than a random distribution. We performed the permutation test with the “randomizeRegion” function and 1000 permutations. Randomizations were performed per chromosome.

We quantified TE impacts on intra-specific small-scale chromosomal rearrangements in the nine isolates of *Z. tritici*. For this, we used the software SyRi (Goel *et al*. 2019) to identify synteny breaks of a minimum of 500bp. We assessed the distances from these synteny breaks to the closest transposons using bedtools “closest” function (Quinlan and Hall 2010).

### Repeat-Induced Point mutation (RIP) analysis

Repeat-Induced Point (RIP) mutation indices were calculated, and Large RIP-affected genomic regions were determined, using the RIPper software (https://github.com/TheRIPper-Fungi/TheRIPper/ (van Wyk *et al*. 2019). Regions of more than 4000 bp that are consecutively affected by RIP are considered to be “large RIP affected genomic regions”. For genome-wide RIP index assessments, we used default parameters of parameters of 1000bp windows with a 500bp step size. RIP Composite index values were calculated as follows: (TpA/ ApT) – (CpA + TpG/ ApC + GpT). A region is affected by RIP when the Composite index is > 0 (van Wyk *et al*. 2019). To calculate the RIP Composite index of each transposon copy, we used 50bp non-overlapping windows using homemade script.

### Data Availability

All data produced and analysed in this study including transposable elements annotations and consensus libraries are available at {ZENODO DOI TO BE ADDED WHEN ACCEPTED}.

## Results

### Transposable elements content varies in the genomes of the *Zymoseptoria* species-complex

Overall, the TE proportions of *Z. passerinii*, *Z. ardabiliae*, *Z. brevis* and *Z. pseudotritici* genomes are higher than the TE proportions of *Z. tritici* isolates (Table S1; Figure 1). Outside of *Z. tritici*, TE content ranges from 6.9 Mb in the *Z. ardabiliae* Za17 genome (18.2% of the genome) to 12.9 Mb in the *Z. passerinii* Zpa63 genome (31.4% of the genome; Figure 1B; Table S1). Among *Z. tritici* isolates, TE content ranges from 5.70 Mb in the Iranian isolate Zt289 genome (14.5% of the genome) to 8.21 Mb in the Zt05 genome (19.8% of the genome) (Figure 1B; Table S1). Most of the TE coverage consists of LTR-retrotransposons among all genomes of *Zymoseptoria* species. LTRs elements represent from 3.2 Mb to 7.4 Mb of the genomes of *Z. tritici* (Zt289) and *Z. passerinii*, respectively. Among these LTRs, we only found the *Copia* and *Gypsy* elements, except in the Iranian *Z. tritici* isolate Zt469, in which we also identified elements belonging to the unique *Bel-Pao* family (representing 0.85 Mb; Table S1). LINE elements are completely absent from the genome of *Z. pseudotritici* and comprise only 0.29% (0.1 Mb) of the genome of *Z. ardabiliae* and 0.5% (0.2 Mb) of the genome of *Z. passerinii*. In contrast, *Z. brevis* Zb87 appears as an outlier: its genome contains 1.2 Mb (2.9% of the genome) of LINE elements. Among *Z. tritici* isolates, LINE content ranges from 0.29% (in Zt469) to 3.83% (in Zt05). Taken together, these results suggest recent invasion and variable expansions of LINEs in *Z. tritici* and *Z. brevis* (Figure 1; Table S1).

**Figure 1:**
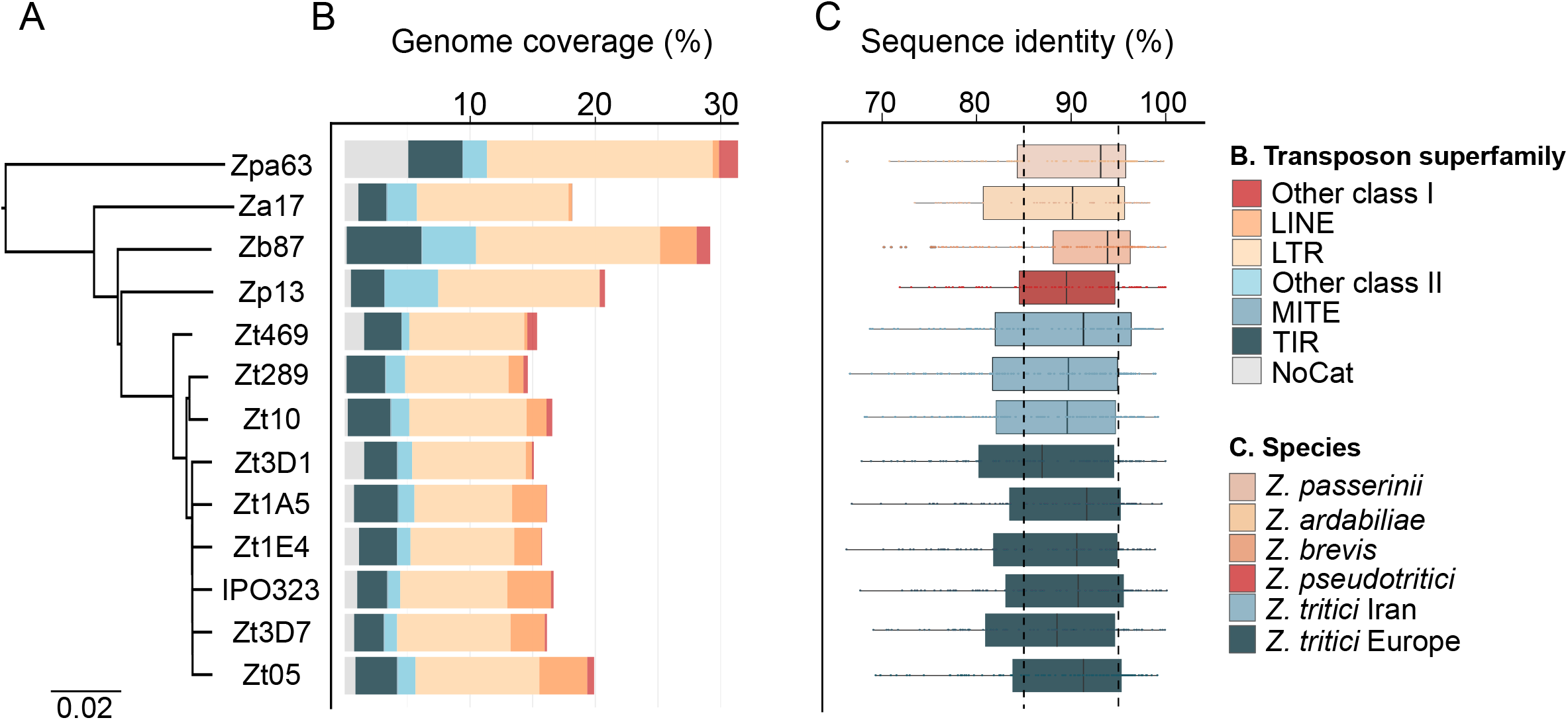
Transposable element content and identity variation in the *Zymoseptoria* genus. A) Phylogenetic tree constructed by the software Andi (Haubold *et al*. 2015), using whole genome assemblies. The tree was rooted with *Cercospora beticola* (Vaghefi *et al*. 2017) as outgroup (data not shown). B) Bars represent TE content (%) per genome estimated after REPET (Flutre *et al*. 2011) annotation. Colours represent TE order coverage with retrotransposons (LTR, LINE and other class I orders in warm colors) and DNA transposons (TIR, MITE and other class II orders in cold colors). C) Sequence identity distribution between TE copies to their cognate consensus. Each dot represents the median sequence identity of TE cluster. Boxplots are coloured in regards to the species and isolate geographical origin.

### Many recent TE insertions postdate diversification of the *Zymoseptoria* genus

One third of the TEs in the different *Zymoseptoria* genomes are relatively recent insertions. Sequence similarities between TEs and their cognate consensus sequences in the genomes of *Zymoseptoria* spp. range from 65.4% to 100% of sequence identity (Figure 1C). Based on the approach used in Maumus & Quesneville (2014), we defined TE categories based on thresholds as follows: 1) copies with less than 85% sequence identity to the consensus comprise *old* insertions and 2) copies with 85% to 95% sequence identity are *intermediate* insertions and 3) copies with more than 95% identity with the consensus sequence represent *recent* insertions. Based on these criteria, we show that *old* insertions represent 33% (*Z. passerinii* – 5.6 Mb) to 46.6% (*Z. pseudotritici* – 5.6 Mb) of the total TE content, while *recent* insertions represent 29.3% (*Z. tritici* Zt3D1 – 3.4Mb) to 40.5% (*Z. brevis* – 6.2Mb) of transposons (Figure 1C; Figure S1). The majority of *recent* TE insertions are retrotransposons while the majority of *old* insertions are retrotransposons and DNA transposons (Figure S2). Taken together these results suggest that at least a fraction of TEs have been recently active within genomes. Variation in TE insertions among *Z. tritici* core chromosomes indicates recent TE transposition activity, mostly driven by LTRs. The proportion of TEs per core chromosome or contig varies from 5.7% (chr11 of Zt469) to 33.3% (chr9 of Zt05) (Figure 2). In addition, isolates with very close TE content per chromosome show very different TE distribution patterns (Figure 2A; Figure S3A). For instance, Zt1A5 and Zt1E4 exhibit the same number of TE insertions on chromosome 9 (163 TEs), however the distribution of these copies is different between the two genomes (Figure 2A). We counted up to three times more TE insertions per chromosome among the nine *Z. tritici* isolates (e.g. 68 vs 198 on chromosome 8) (Figure 2B). Variation is mostly driven by LTR retrotransposon content (Figure S3B). Taken together these results indicate that each *Z. tritici* genome reflects independent TE insertion events and the recent transposition activity of few specific TE orders (e.g. LTRs).

**Figure 2:**
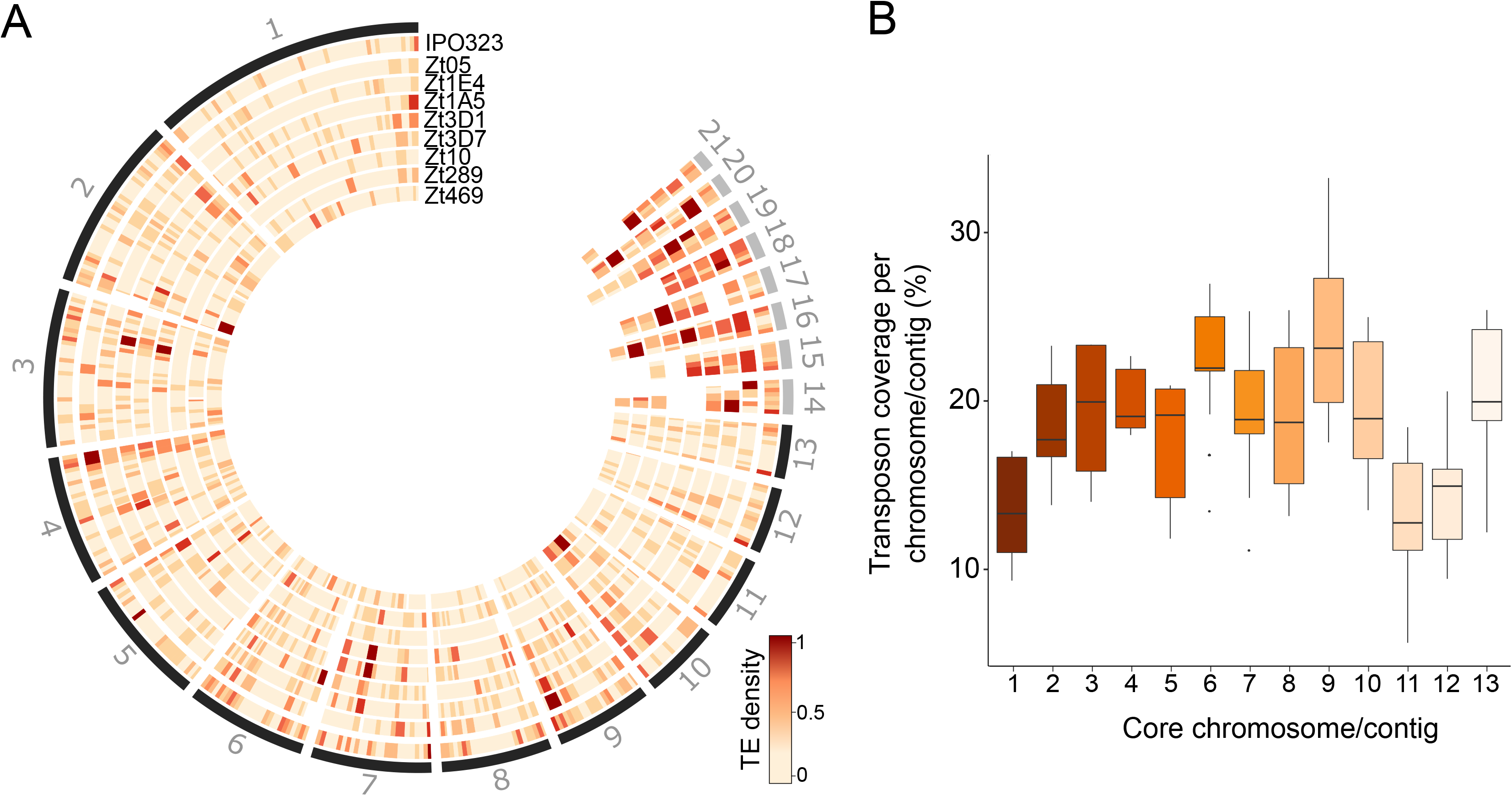
Transposable element content variation along chromosomes of nine *Z. tritici* isolates. A) Circos plot of the TE content along chromosomes of nine *Z. tritici* isolates. The first track represents the karyotype of the reference isolate IPO323; the second track shows a heatmap of TE density per 100kb windows in IPO323. Track three to ten represent the density of TE copies per 100kb windows in other *Z. tritici* isolates. Chromosomal coordinates refer to the closest orthologous gene projected on the IPO323 genome. Darker colors correspond to a higher TE density. B) TE content variation of the 13 core chromosomes of *Z. tritici*. The boxplots represent the distribution of TE coverage percentage per chromosome or contig among the nine *Z. tritici* isolates.

### Past and present TE activity has shaped the genomes of *Zymoseptoria* species

TE-rich accessory chromosomes represent an ancestral trait of the genome architecture among the *Zymoseptoria* species-complex. In agreement with a previous study (Grandaubert *et al*. 2015), transposons in this study are enriched on accessory chromosomes (Figure 3A). Rearrangements in the genomes also co-occur in the vicinity of TEs, indicating the TEs’ potential role in causing structural variation among *Zymoseptoria* genomes. We identified a large interspecies chromosomal inversion on contig_65 of *Z. brevis* Zb87 and contig_76 of *Z. pseudotritici* Zp13 compared to the chromosome 2 of *Z. tritici* IPO323 (Feurtey *et al*. 2019). This large rearranging region actually consists of several inversions between clusters of TEs (Figure S4). We identified three TE clusters in Zb87 (46 TEs representing 0.12 Mb) and Zp13 (30 TEs representing 0.28 Mb) that surround the inverted loci and are absent in *Z. tritici* IPO323 (Figure S4). We further scrutinized the link between TEs and rearrangements by pairwise comparisons of the nine *Z. tritici* isolates to the reference IPO323 (Table S3). Of all inverted regions identified per isolate, 27% (Zt469) to 72% (Zt1A5) are flanked (separated by less than 10bp) by transposons both upstream and downstream (Table S3). In total, we counted from 15 (Zt10) to 52 (Zt1E4) intra-specific inversions comprising more than 500 bp in pairwise comparisons to the reference isolate IPO323 (Table S3). The cumulative size of these inversions ranges from ~136.7 kb to ~845.2 kb in Zt1E4 and Zt469, respectively. From 10 (Zt469) to 128 (Zt05) genes are located in inverted genomic regions (Table S3). One inverted region of 170kb is found on chromosome 13 in the reference genome but absent from the other isolates. Taken together these results indicate that TEs have shaped genome rearrangements and genome compartmentalization in *Zymoseptoria* species.

**Figure 3:**
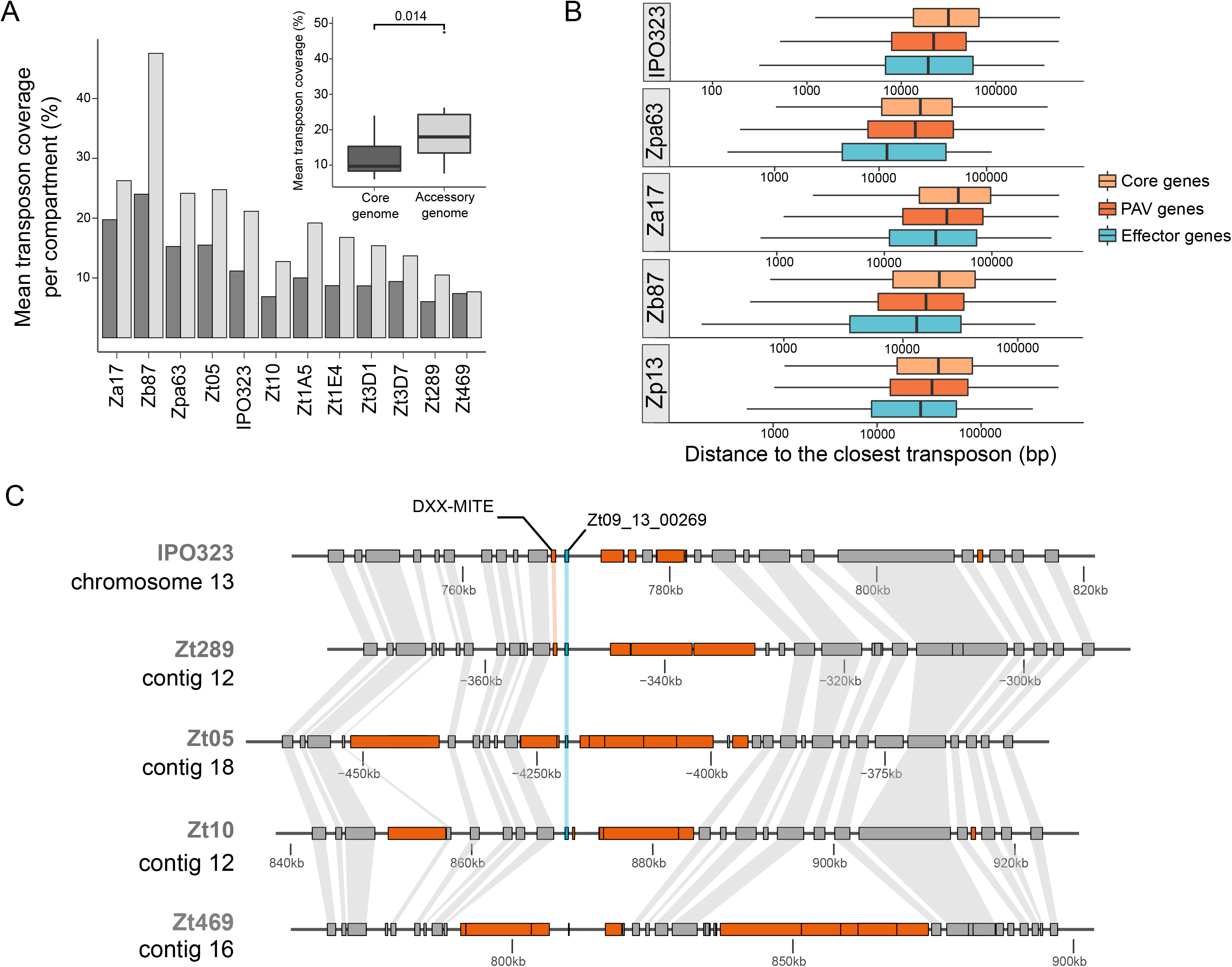
Transposable element insertions shape genome compartmentalization and plasticity. A) Bars represents mean percentage of TE content per genome compartment of *Zymoseptoria* species of core (dark grey) and non-core (light grey) chromosomes/contigs. Boxplots describe the mean TE coverage percentage per genome compartment among all thirteen genomes of *Zymoseptoria* species. P-value was estimated using a Kruskal-Wallis test. B) Distribution of distance between effectors (blue), species- (red) and genus-specific (dark orange) genes and core genes (light orange) to the closest TE per chromosome/contig. Mean distances and permutation test results per gene category are summarised in Table S3. C) Example of a presence/absence polymorphism of the candidate effector Zt09_13_00269 identified in (Haueisen *et al*. 2018) in the vicinity of TE-rich region. TEs are shown in orange, genes in grey and the effector of interest is shown in blue boxes. Connecting lines in grey represent the orthologous genes in each genome.

### Presence/absence variation and effector genes associate with transposable elements

Presence/absence variable (PAV) genes are enriched in the vicinity of TEs while core genes are located further apart than expected by chance. To assess the potential impact of TEs on the high gene presence/absence variation described in *Z. tritici* (Plissonneau *et al*. 2018; Badet *et al*. 2019), we explored the genes close to TEs. We found that PAV genes are significantly closer to TEs in the genomes of *Z. ardabiliae, Z. brevis*, and the three *Z. tritici* isolates IPO323, Zt05 and Zt10 (permutation test of 1000 iterations, p-value< 0.05) (Figure 3B; Table S4). On the contrary, the distance between core genes and TEs is significantly higher than expected from a random distribution (permutation test of 1000 iterations p-value< 0.05). The exceptions are in *Z. pseudotritici*, and *Z. passerinii* in which observed distances between PAV genes and core genes to TEs were not significantly different from a random distribution (Table S4).

To assess if particular TE families are associated with genes, we tested the distribution of the major TE families from class I (i.e. *Copia, Gypsy* and *LINE* elements) and class II (*TIR, Helitron* and *MITE* elements) (Table S5). In all *Zymoseptoria* genomes *TIR* and *MITE* elements are located significantly closer to PAV genes than expected from a random distribution (applying a permutation test of 1000 iteration p-value< 0.05) (Table S5). Also *Copia* elements are significantly closer to PAV genes than expected from a random distribution in *Z. tritici* (IPO323 and Zt10) and *Z. ardabiliae*. In contrast, class I *Gypsy* elements are more distant to PAV genes than expected from a random distribution in all *Zymoseptoria* genomes even though they are the most abundant TEs (permutation test of 1000 iteration p-value< 0.05) (Table S5). In conclusion, different TE families are found closer to PAV genes among the genomes of the *Zymoseptoria* species-complex.

Effector genes are enriched in the vicinity of TEs in the genomes of the *Zymoseptoria* species-complex. We used the previously annotated effector genes of *Z. ardabiliae*, *Z. brevis*, *Z. pseudotritici*, *Z. passerinii* and the three *Z. tritici* isolates IPO323, Zt05 and Zt10 (Feurtey *et al*. 2019) and tested whether they are enriched in the vicinity of TEs. We show that these effector genes are significantly closer to TEs than expected if they were randomly distributed in *Zymoseptoria* genomes (permutation test of 1000 iteration p-value< 0.05) (Figure 3B; Table S4). Effector genes associate with TEs in each genome, but not with the same elements. Effector genes of *Z. tritici* IPO323 associate with *TIR* elements while in Zt10 effector genes associate with *Helitron* and *Copia* elements but not *TIR* elements (Table S5). As an example, the effector gene Zt09_13_00269 shows a presence/absence polymorphism among *Zymoseptoria tritici* isolates (Figure 3C). This effector is absent from the *Aegilops*-infecting isolate Zt469 which could indicate a potential link with host specificity. Zt09_13_00269 is surrounded by different TEs both up- and downstream. Interestingly, this effector candidate is expressed during wheat infection in the reference isolate IPO323 (56.9 transcripts per million – TPM), while not expressed in Zt05 and Zt10 during wheat infection (1.23 TPM and 4.13 TPM respectively) (Table S6). A specific MITE transposon localizes 881 bp upstream of Zt09_13_00269 in IPO323 and in Zt289 but not in the other isolates (Figure 3C). In conclusion, effectors genes are enriched in the vicinity of TEs suggesting that TEs could play a role in the evolution of pathogenicity-related genes in the *Zymoseptoria* species-complex.

### Signatures of Repeat-Induced Point mutations are reduced on TEs of European *Z. tritici* isolates compared to other *Zymoseptoria* genomes

RIP efficacy recently decreased in European *Z. tritici* isolates. We here evaluated the RIP signatures in the *Zymoseptoria* genomes to estimate to what extent RIP affects different species of the genus. To this end, we scanned each genome and calculated RIP indices (see Methods). In total, RIP signatures affect between 17.4% (in Zt3D7) and 34.5% (in Zpa63) of the total genomic TE content (Table 1). These RIP proportions correspond to the TE content of each genome indicating RIP has been – at least recently – active in the genomes of *Zymoseptoria* species. Therefore, as the TE content in the genomes of *Zymoseptoria* spp. are recent insertions mostly and RIP is found also on recent copies, it indicates that the RIP has been recently active in the genomes. RIP efficacy is not equivalent for all types of TEs (van Wyk *et al*. 2019). The small-sized and non-autonomous TEs such as MITE DNA-transposons are less affected by RIP (Figure S7). Overall, in *Zymoseptoria*, the vast majority of TEs carries RIP signatures (86-99% of TEs are RIPed), based on the estimation of RIP composite indices per 50 bp windows of TEs (Figure 4). The large number of RIPed TEs suggest a RIP-induced decay of TE copies in the genomes of *Zymoseptoria* species. However, 8.1% to 13.5% of TEs in the European *Z. tritici* isolates do not have any RIP signature, while this percentage is only of 1 to 6.8% for TEs of the Iranian *Z. tritici* isolates (Figure 4; Figure S6). This reduction of RIP can be linked to the variation in TE repertoires among the genomes of *Z. tritici* isolates. We observed that the proportion of MITE elements (i.e. elements that are less affected by RIP) do not correlate with this reduction of RIP in copies. The Iranian *Z. tritici* isolates comprise from 29 to 121 MITE copies while European isolates comprise from 68 to 156 MITE copies (Table S1). The reduction of RIP signatures in European *Z. tritici* isolates indicates a relaxation of the RIP efficacy in those isolates, potentially due to a different composition in TE repertoires.

**Table 1:**
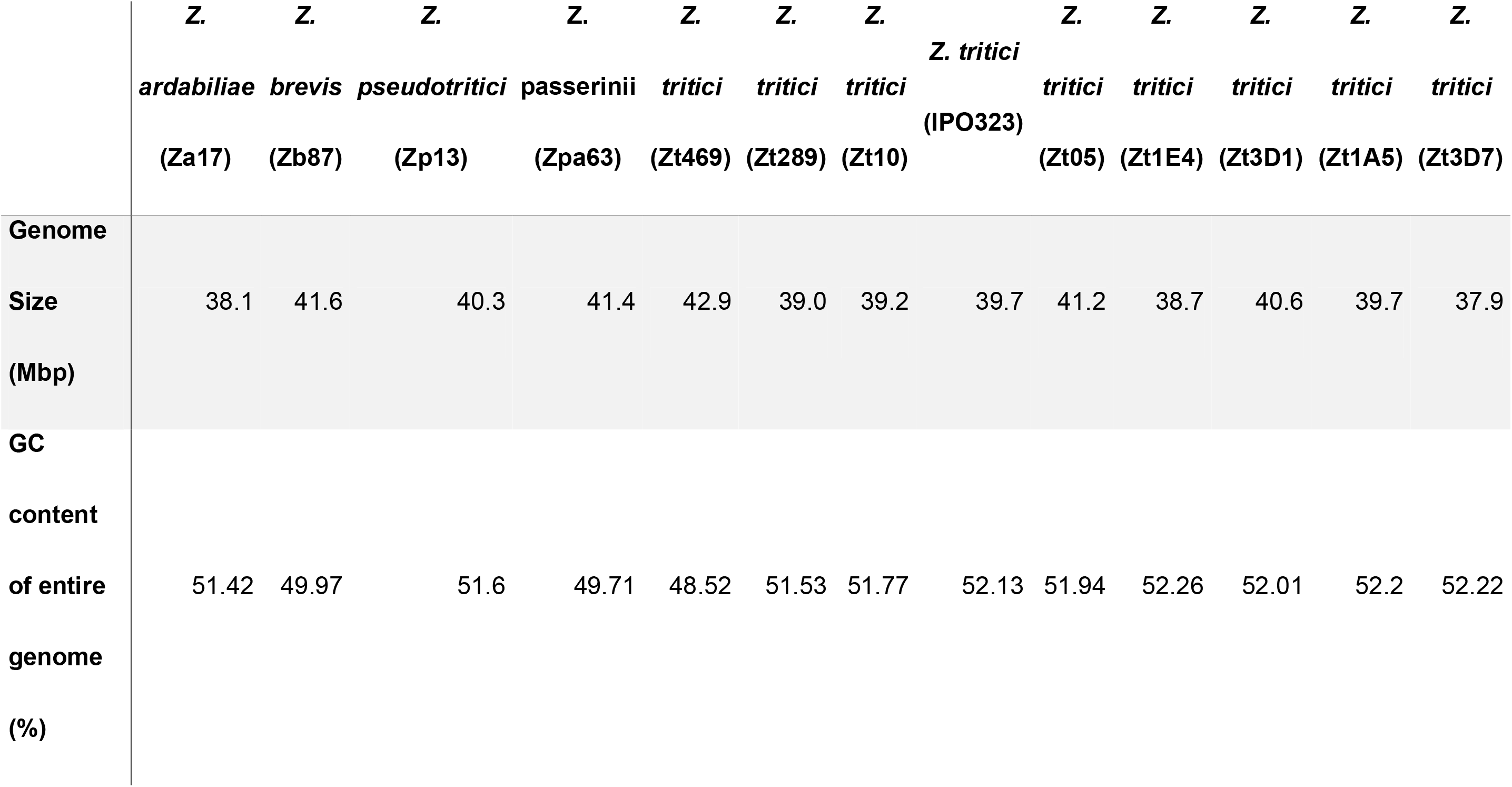
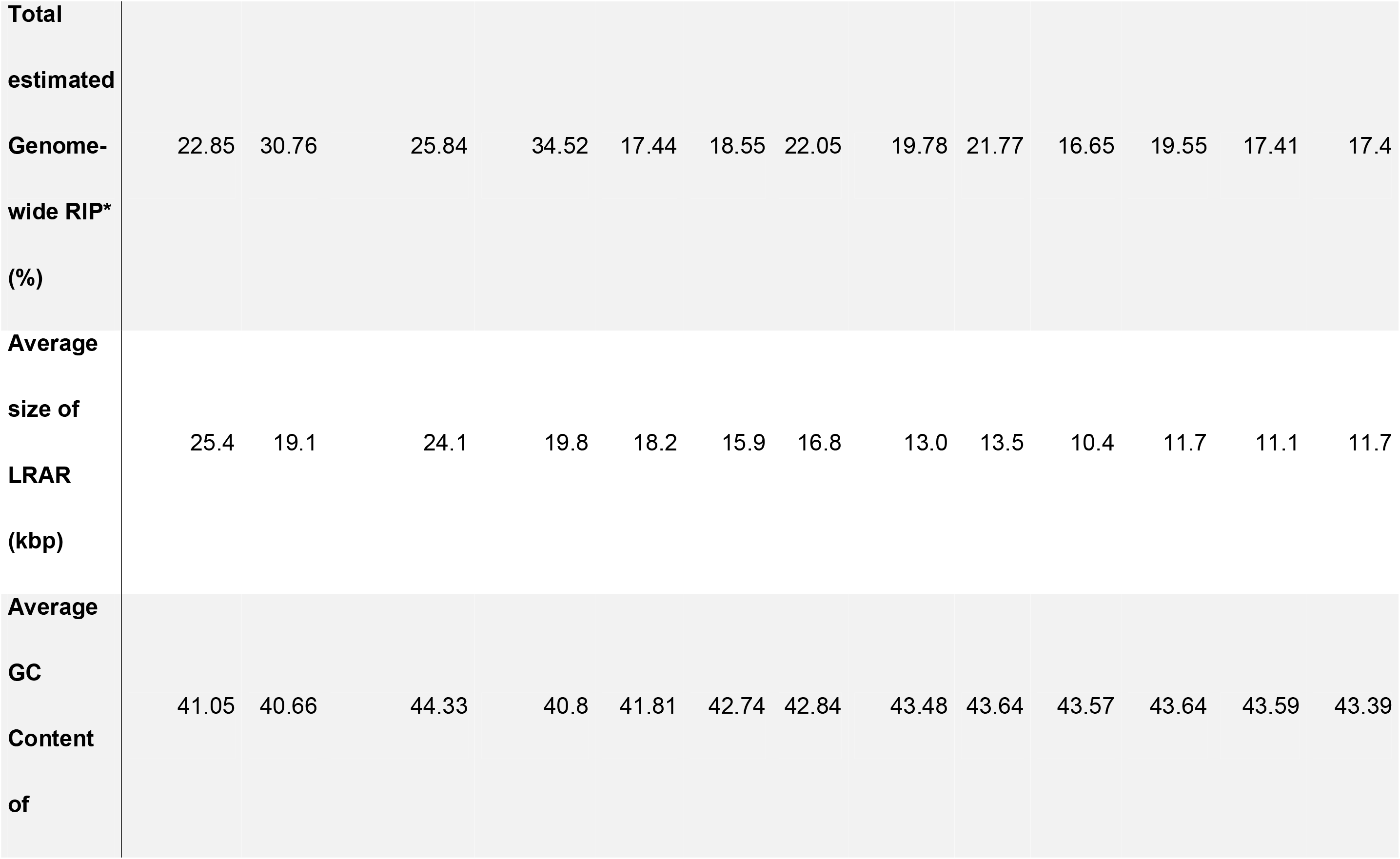
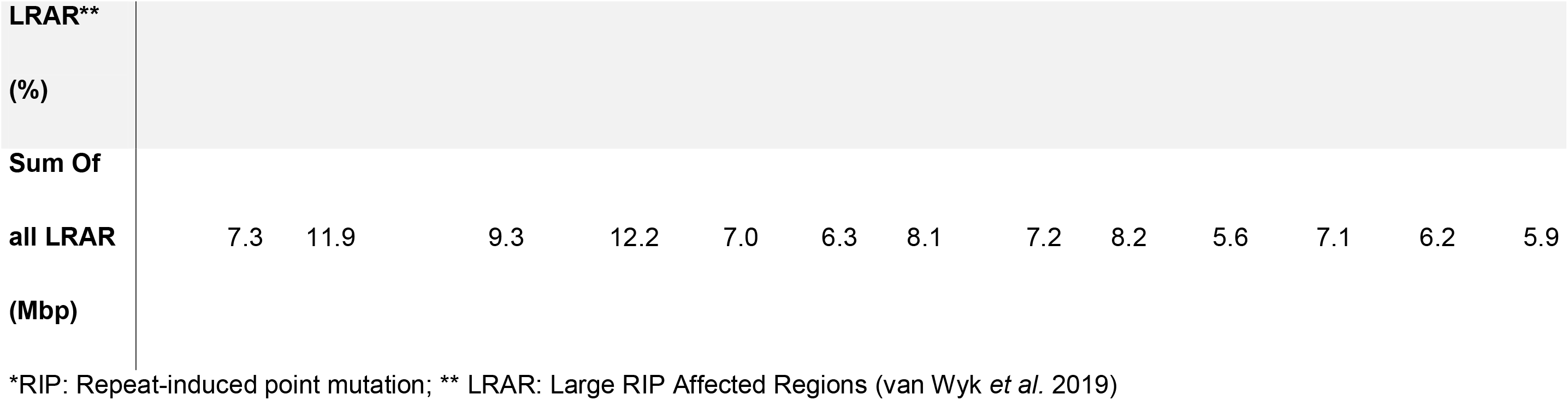
Repeat-induced point mutation signatures in the genomes of *Zymoseptoria* species

**Figure 4:**
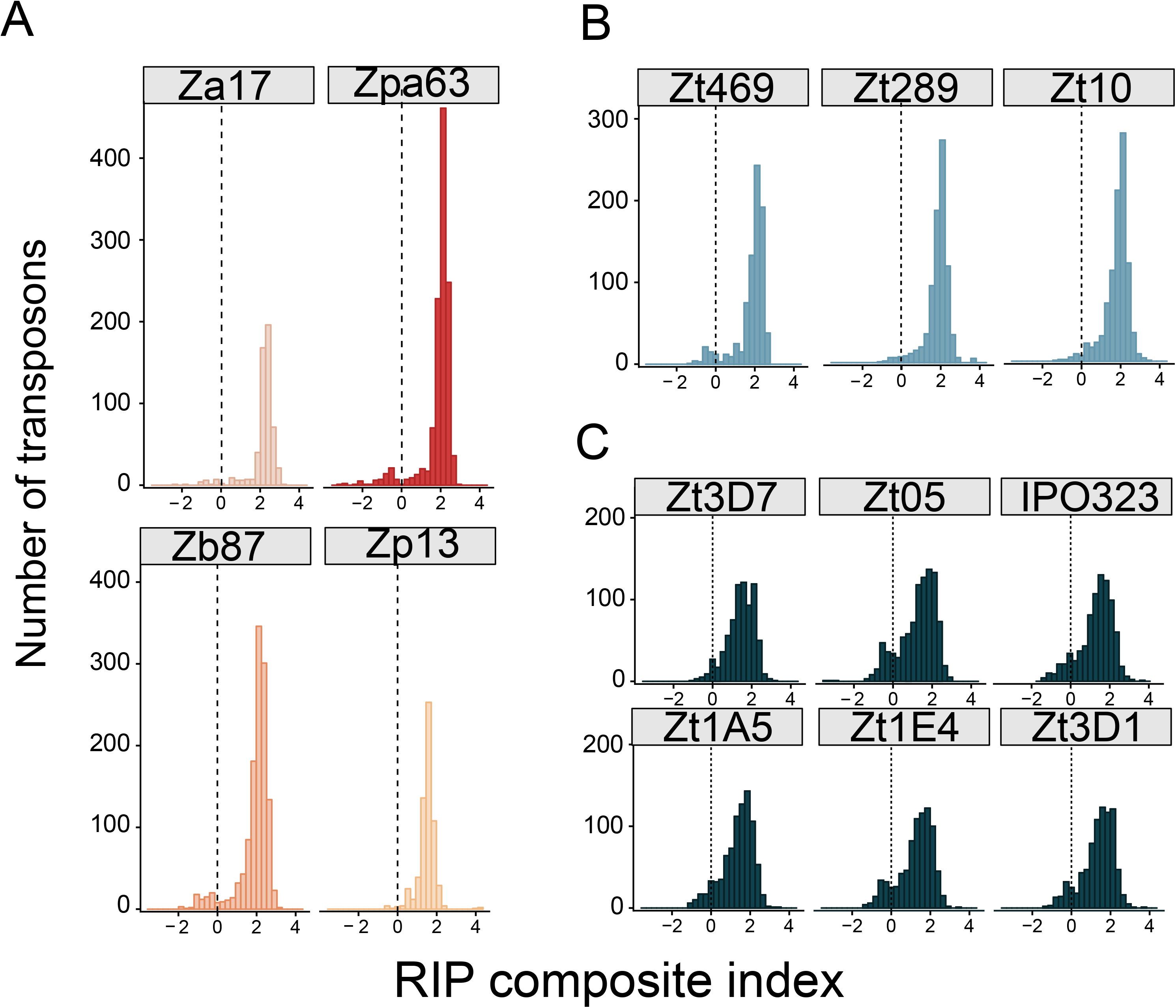
Repeat Induced Point (RIP) mutations in transposable elements of *Zymoseptoria* spp. genomes. Histograms of Composite RIP index (CRI) frequencies of TEs estimated using a 50bp sliding windows approach as follows: CRI = (TpA/ ApT) – (CpA + TpG/ ApC + GpT) for A) *Z*. *passerinii*, *Z. ardabiliae, Z. brevis* and *Z. pseudotritici*, and B) Iranian *Z. tritici* isolates and C) European *Z. tritici* isolates. Vertical dash lines exhibit the threshold (0) above which CRI values indicate a RIP signature.

In addition, the average size of Large RIP-affected regions (LRARs) can be used as a proxy for RIP mechanism efficiency because it reflects to which extent a highly repeated region is affected by RIP. In the genomes of *Zymoseptoria* species, LRARs have an average size between 10.3 kb (in Zt1E4) and 25.4 kb (in Za17) and comprise large AT-rich regions (Table 1). LRAR average sizes in the *Z. tritici* European isolates are reduced compared to the other members of the *Zymoseptoria* species-complex (Table 1). It is worth noting that *Z. tritici* genomes with lower LRARs average size do not have significantly lower TE content compared to the Iranian isolates. This indicate that the average size of LRARs is a good indicator for RIP efficiency. Besides TE sequences, only a small number of genes exhibited RIP signatures, including 43 genes in *Z. pseudotritici* and 92 genes in *Z. tritici* Zt05 (Table S7). Taken together, we conclude that RIP is highly efficient to inactivate TEs in the genomes of *Zymoseptoria* species but show evidence of relaxation in the genomes of the European isolates of *Z. tritici*.

## Discussion

Growing evidence demonstrates that transposable elements represent key players for the evolution and adaptation of fungal plant pathogens (Möller and Stukenbrock 2017). We investigated how past and present transposition events have shaped genome evolution of a major wheat pathogen and its wild-grass infecting sister-species. Within the genus *Zymoseptoria*, TEs associate with effector genes and PAV genes, as previously described for isolates of *Z. tritici* (Plissonneau *et al*. 2018). This suggests that TEs are either directly or indirectly involved in evolution of effectors and PAV genes in *Zymoseptoria* species (Faino *et al*. 2016). TEs may therefore represent a major driver of *Zymoseptoria* genome evolution.

Our detailed transposon analysis confirmed a conserved genome compartmentalization of TEs in accessory regions among species of the *Zymoseptoria* genus. TE accumulation is often associated with genome size expansions (Raffaele and Kamoun 2012). However, genome sizes of *Zymoseptoria* spp. are comparatively stable; even a duplication of the transposon content does not cause genome size variation larger than 2 Mb (Feurtey *et al*. 2019). This can be explained by the high content of accessory chromosomes showing presence/absence variation: indeed, these chromosomes are small but have a high content of TEs (Grandaubert *et al*. 2015). This was previously shown for *Z. tritici*, and here we show that genome compartmentalization involving dynamic TE content represents a more ancestral trait of the genomes of these fungi (Goodwin *et al*. 2011). Purifying selection pressure is higher on the gene-dense core chromosomes of *Z. tritici* compared to the transposon-dense accessory chromosomes (Grandaubert *et al*. 2019). It is possible that relaxed selection acting on accessory genome compartments allows TEs to accumulate in these regions.

TEs are involved either directly or indirectly in the genome plasticity of fungi. In this study, we explored the impact of TEs on genome architecture and gene presence/absence variation between and within species of *Zymoseptoria*. TEs associate with regions exhibiting inter- and intra-species chromosomal rearrangements. Furthermore, we found that genes affected by presence/absence variation between and within species are located close to TEs. Such association of gene presence/absence with TEs has also been observed in other plant pathogenic fungi such as the rice blast *M. oryzae* (Yoshida *et al*. 2016; Bao *et al*. 2017).

TEs associate with effector genes in *Zymoseptoria* species, supporting the importance of TE-driven gene evolution for pathogenicity. In the wheat pathogen *Pyrenophora tritici-repentis*, the pathogenicity-related protein ToxA, is found in the two other wheat-associated pathogens *Parastagonospora nodorum* and *Bipolaris sorokiniana* (McDonald *et al*. 2019). Horizontal transfer of the ToxA encoding gene was demonstrated between these wheat-associated species and transfer was mediated by a DNA transposon from the hAT family (McDonald *et al*. 2019). TE insertions that facilitate variation of pathogenicity-related traits have been reported in several other fungal pathogens, including *L. maculans*, *F. oxysporum*, *M. oryzae* and *Verticillium dahliae* (Rouxel *et al*. 2011; Amselem *et al*. 2015a; Faino *et al*. 2016; Fokkens *et al*. 2018).

The majority of TEs in the genomes of *Zymoseptoria* species represents young insertions. TE insertions in these closely related species are species- or strain-specific and one third comprise copies of highly conserved sequences. Extensive copy number variation along core chromosomes in the nine *Z. tritici* isolates strengthens our conclusion that the majority of TEs has spread recently. A study of TE dynamics in worldwide *Z. tritici* populations demonstrated expansions of TE families in isolates of recently founded populations compared to isolates collected close to the center of origin (Oggenfuss *et al*. 2020). More investigations are needed to understand how TE expansions in *Z. tritici* closely related species occurred, particularly for the higher repeated genomes of *Z. passerinii* and *Z. brevis*. It would be interesting to explore if TEs also accumulate more in genomes of *Z. passerinii* and *Z. brevis* isolates outside of the center of origins or if populations show different TE accumulation patterns. Understanding how TEs have accumulated stronger in these genomes requires comparing more *Z. passerinii* and *Z. brevis* isolates.

The high level of young TE insertions in *Zymoseptoria* species contrasts with the high level of RIP mutations found in TE copies. The classical model of TE burst and decay dynamics states that TE proliferation is counterbalanced by genome defence mechanisms with more or less efficacy, which eventually leads to TE elimination (Arkhipova 2018). Here, we observe high level of RIP signatures even on recent TE copies suggesting that most TEs are likely inactive. Based on this, we speculate that the majority of TEs in *Zymoseptoria* were affected by RIP shortly after their insertion. It is worth noting that inactive TEs can still spread in genomes to a certain extent. Indeed, recombination between homologous regions can lead to duplications (Bourque *et al*. 2018).

The highly efficient RIP system may impede evolutionary innovation among *Zymoseptoria*. RIP prevents gene duplications in addition to TE duplication, and gene duplications are considered a major driver of genome evolution (Galagan and Selker 2004). Although RIP can be ‘leaky’ and induce mutation on single-copy genes, which has been demonstrated as efficient mechanism for effector diversification in some fungal species (Rouxel *et al*. 2011). We found however very few genes harbouring RIP signatures among the genomes of the *Zymoseptoria* species, which could indicate that such mechanism of effector diversification via RIP scarcely occur in *Zymoseptoria* species. The details of how RIP mechanism occurs in *Zymoseptoria* spp. remain largely unknown. However, one key player in RIP is the DNA methyltransferase DIM-2. RIP reduction in European *Z. tritici* isolates correlates with a deficiency in DNA methylation and absence of DIM-2 proteins in European *Z. tritici* isolates (Dhillon *et al*. 2010; Möller *et al*. 2020). Host genome defences against TEs represent a potential fitness cost notably with regard to gene duplication for rapid adaptation (Galagan and Selker 2004). For example, there are almost no gene duplications in the genome of the model species *N. crassa* because of extremely high RIP efficacy (Galagan and Selker 2004). In *Z. tritici*, a recent study demonstrated the TE-induced expansion of MSF transporters (Oggenfuss *et al*. 2020). The latter are mostly involved in fungicide resistance in *Z. tritici* (Omrane *et al*. 2017a).

Moderate transposon activity has the potential to be advantageous for rapid adaptation of fungal plant-pathogens. We propose that the relaxation of transposition repression by RIP in *Z. tritici* isolates outside of the center of origin could represent an advantage for rapid adaptation. For example, the increase of TE insertions in *Z. tritici* could be advantageous during migration to new environments following wheat deployment across the world (Oggenfuss *et al*. 2020). In the cereal pathogen *M. oryzae*, DIM-2 is functional in isolates infecting wheat, rice and common millet, but isolates infecting foxtail millet carry a non-functional DIM-2 variant. There may be a link between DNA methylation of transposons and infection success of *M. oryzae* on specific host plant species (Ikeda *et al*. 2013). Based on the findings of our study, relaxation in RIP efficacy among *Z. tritici* isolates might potentially be advantageous for rapid adaptation to new wheat cultivars.

## Acknowledgements

CL was funded by the Institut national de la recherche agronomique (INRA) in the framework of a “Contrat Jeune Scientifique” and by the Labex ARBRE (Lab of Excellence ARBRE). Genome research in the group of EHS is funded by the Max Planck Society and CIFAR. AF is funded by the DFG Priority Program SPP1819. CL performed data and results analyses. EHS and CL contributed to the design and implementation of the research. All authors contributed to the writing of the manuscript. All authors read and approved the manuscript.

---

**Figure S1: Burst and decay TE evolution.** Insertion of a TE copy (brown box) at a locus of the host genome, followed by further insertions during time and an accumulation of mutations (dark red triangles) and structural modifications such as partial deletions (dash lines) and/or insertions of other TEs (gold box). The older the insertion the more variants it accumulates. Therefore, TE families with old insertions are less similar to their consensus sequence while younger insertions are highly similar to the consensus. Consensus sequence represents the ancestral sequence of each annotated TE cluster (Flutre *et al*. 2011).

**Figure S2: Relative abundance of TE divergent (left panel) and conserved (right panel) copies per genome.** The divergent copies have less than 85% of sequence identity to their cognate consensus while conserved copies have more than 95% of sequence identity to the consensus. Retrotransposons are represented in warm colors (LTR in light orange, LINE in orange and other retrotransposons orders in red) and DNA transposons are represented in cold colors (TIR in dark blue, MITE in blue and other DNA transposons in light blue). Uncategorized TEs are represented in grey.

**Figure S3: Intraspecific variation in TE coverage per chromosome/contig** A) Total TE coverage per core chromosome/contig in the genomes of *Z. tritici* isolates. B) TE orders coverage contributions per core chromosome/contig of nine *Z.tritici* isolates.

**Figure S4: Chromosomal inversion identified in *Z. tritici* IPO323 compared to *Z. brevis* Zb87 and *Z. pseudotritici* Zp13.** TEs are represented in orange, genes in grey and effectors genes in blue. Links indicate orthologous genes. Arrows represent effector genes loci.

**Figure S5: Comparison of Repeat-induced point mutation (RIP) composite index variation per sample group.** Distribution comparison of RIP index (CRI) frequencies of TEs estimated using a 50bp sliding windows approach as follows: CRI = (TpA/ ApT) – (CpA + TpG/ ApC + GpT) for Sister species: *Z*. *passerinii, Z. ardabiliae, Z. brevis* and *Z. pseudotritici* (light orange), Iranian *Z. tritici* isolates (light blue) and C) European *Z. tritici* isolates (dark blue). P-values were estimated using Kruskal-Wallis test.

**Figure S6: Repeat-induced point mutation (RIP) composite index per TE copy of all *Zymoseptoria* species.** Mean RIP composite index (CRI) frequencies of TEs estimated using a 50bp sliding windows approach as follows: CRI = (TpA/ ApT) – (CpA + TpG/ ApC + GpT) for each TE copy per order from Class I and Class II.

**Table S1: TE content in thirteen genomes of *Zymoseptoria* genus metrics.** TE classification was performed based on Wicker’s classification system (Wicker *et al*. 2007). The two classes (Class I and Class II) of TE are subdivided into subclasses, orders and superfamilies as follows: Long terminal repeats (LTR) elements, *Dictyostelium* intermediate repeat sequence (DIRS), Penelope-like elements (PLEs), Long INterspersed Elements (LINEs) and Short INterspersed Elements (SINEs). Terminal inverted repeat (TIR), Crypton, Helitron, Maverick and Miniature inverted-repeat transposable element (MITEs).

**Table S2: Metrics output from REPET annotation of TEs.**

**Table S3: Chromosomal inversions between *Z. tritici* isolates compared to the reference IPO323.**

**Table S4: Permutations tests based on mean distance of TEs to genes.**

**TableS5: Permutations tests based on mean distance of TE orders to genes.**

**TableS6: Effectors candidates and their neighbouring TEs**

**Table S7: Genes affected by Repeat-induced point mutations signatures**

